# Silkworm model for *Bacillus anthracis* infection and virulence determination

**DOI:** 10.1101/2021.06.12.448172

**Authors:** Atmika Paudel, Yoshikazu Furuta, Hideaki Higashi

**Affiliations:** Division of Infection and Immunity, International Institute for Zoonosis Control, Hokkaido University, Sapporo, Hokkaido, Japan

**Keywords:** *Bacillus anthracis*, *Bombyx mori*, silkworm, virulence, animal model, host-pathogen interaction

## Abstract

*Bacillus anthracis* is an obligate pathogen and a causative agent of anthrax. Its major virulence factors are plasmid-coded; however, recent studies have revealed chromosome-encoded virulence factors, indicating that the current understanding of its virulence mechanism is elusive and needs further investigation. In this study, we established a silkworm (*Bombyx mori*) infection model of *B. anthracis* Sterne. We showed that silkworms were killed by *B. anthracis* and cured of the infection when administered with antibiotics. We quantitatively determined the lethal dose of the bacteria that kills 50% larvae and effective doses of antibiotics that cure 50% infected larvae. Furthermore, we demonstrated that *B. anthracis* mutants with disruption in virulence genes such as *pagA, lef*, and *atxA* had attenuated silkworm-killing ability and reduced colonization in silkworm hemolymph. The silkworm infection model established in this study can be utilized in large-scale infection experiments to identify novel virulence determinants and develop novel therapeutic options against *B. anthracis* infections.

## Introduction

*Bacillus anthracis* is a spore-forming Gram-positive bacterium that infects both animals and humans. Most animals ingest *B. anthracis* spores while grazing and develop an infection, and humans occasionally acquire infection from the infected animals or animal products. The spore-forming ability allows the bacteria to exist for decades in a dormant state and resist harsh environments^1^. Despite immunization efforts, *B. anthracis* is still a potential threat due to sporadic anthrax outbreaks^2–9^ and its use in bioterrorism^10^. Consequently, the world health organization and centers for disease control and prevention have placed *B. anthracis* as one of the top bioterrorism agents.

The pathogenicity of *B. anthracis* has been attributed mainly to toxins and capsule encoded in the plasmids pXO1 and pXO2, respectively^11,12^. Emerging evidence has uncovered the involvement of chromosomal genes in virulence^13–18^, implying that other virulence factors of *B. anthracis* are yet to be identified. As pathogenesis is an outcome of host-pathogen interaction, a suitable animal host is desired to understand the virulence mechanism and design novel therapeutic approaches. A model that can be used in large numbers and have fewer ethical concerns would be of particular importance during the initial phases of research. Due to economic, technical, and ethical concerns associated with the use of vertebrate animals, scientists are turning toward invertebrate animal models such as *Caenorhabditis elegans*^19–21^, *Drosophila melanogaster*^22–24^, *Galleria mellonella*^25–27^ and *Bombyx mori*^23–30^ for large scale screenings. While accurate dose administration in *C. elegans* and *D. melanogaster* is difficult due to their small size, the use of *G. mellonella* is challenging due to its faster locomotion. *Bombyx mori* (silkworm) larvae have been used as a desirable animal model with many physical and biological advantages^31^. They are small enough for easy handling yet large to perform experiments involving organ isolations and desired quantity injections. Besides, due to their slow locomotion and harmless nature, they have less biohazard potential. Conserved basic biological features with mammals make silkworms appropriate as animal models of human diseases^32–34^. Using silkworms as an infection model, novel antimicrobial agents and novel genes with roles in bacterial virulence have been identified^28,29,35–38^.

In this study, we established a silkworm model of *B. anthracis* infection. We demonstrated that *B. anthracis* kills silkworms, and the infection can be treated by clinically used antibiotics. Using fluorescence protein-expressing strain, we revealed how *B. anthracis* establishes infection inside the host. We further showed that mutants with disruption in the genes encoding known virulence factors had decreased virulence in silkworms.

## Materials and Methods

### Bacterial strains and culture condition

The bacterial strains used in this study are shown in **Table 1**. *B. anthracis* 34F2 with pRP1099, a plasmid possessing the gene for AmCyan1 protein, was constructed by conjugation^39^. Strains were grown in Brain-Heart Infusion (BHI) medium (Difco, USA) for routine culture at 37°C. Kanamycin (20 μg/ml) was supplemented for BYF10124. For liquid cultures, strains were grown at 37°C with shaking at 155 rpm.

**Table 1:** Bacterial strains used in this study.

| Strain | Description | Reference |
| --- | --- | --- |
| 34F2 | *Bacillus anthracis* Sterne, vaccine strain (pXO1<sup>+</sup>, pXO2<sup>-</sup>) | Sterne *et al.*<sup>40</sup> |
| BYF10008 | Strain derived from 34F2, $\Delta pagA$ | Furuta *et al.*<sup>41</sup> |
| BYF10054 | Strain derived from 34F2, $\Delta atxA$ | Furuta *et al.*<sup>41</sup> |
| BYF10009 | Strain derived from 34F2, $\Delta lef$ | Furuta *et al.*<sup>41</sup> |
| BYF10124 | 34F2 harboring pRP1099, expressing fluorescence protein AmCyan1 | This study |

### Silkworm rearing

Silkworm eggs were purchased from Ehime-Sanshu Co., Ltd. (Ehime, Japan), disinfected, and reared at 27°C. The worms were fed with antibiotic-containing artificial diet Silkmate 2S (Nihon Nosan Corp., Japan) until the fifth instar stage as previously described^36^. After the larvae turn to the fifth instar, they were fed an antibiotic-free artificial diet (Sysmex, Japan) and used for infection experiments the following day.

### Silkworm infection experiments

For infection experiments, fifth-instar day-2 silkworm larvae were used. Bacterial strains were revived from glycerol stock by streaking them on BHI agar plates and incubating overnight at 37°C. The overnight grown single colony was inoculated and cultured overnight in 5 ml BHI medium with shaking at 155 rpm. The culture was 100-fold diluted in BHI medium and grown till the OD_600_ reached 0.5. The cells were diluted with physiological saline solution (0.9% NaCl), where applicable, and 50 μl of bacterial suspension was injected into the hemolymph of each larva using a 1-ml syringe equipped with a 27-gauze needle (Terumo Medical Corporation, Japan). Silkworms were considered dead if they did not move when poked with forceps.

### Antimicrobial susceptibility test

Antibiotics were obtained from either Fujifilm Wako, Japan, or Sigma Aldrich, Japan. Antimicrobial susceptibility test was performed by broth micro-dilution assay according to the Clinical and Laboratory Standards Institute protocol (CLSI) as explained previously^36^. The plate was incubated at 37°C for 20 h, and the minimum inhibitory concentration (MIC) of each antibiotic was determined as the minimum concentration that inhibited the growth of bacteria.

### Treatment of infection by antibiotics in silkworms

To evaluate the therapeutic activities of clinically used antibiotics in the infected silkworms, exponentially growing bacteria (~5 x 10^2^ CFU) was injected into the hemolymph of each larva. Different concentration of antibiotics was prepared in saline and injected to the larvae into the hemolymph within 30 min of infection. For survival assay, 1 mg/kg of doxycycline and ampicillin each were injected into the larvae. To determine effective doses that cure 50% larvae (ED_50_), various concentrations of doxycycline and ampicillin were prepared in saline and injected into the hemolymph of infected larvae (n=3 for each dose). The survival of larvae was recorded, and ED_50_ was calculated from the survival at 16h post-infection by logistic regression analysis using the logit link function. To determine the microbial burden, larvae infected with *B. anthracis* (6 x 10^2^ CFU/larva) were injected with doxycycline and ampicillin (1 mg/kg; n=10) within 30 min of infection, hemolymph was collected 6h and 9h post-injection, diluted with saline and spread on Luria-Bertani agar plates followed by overnight incubation at 37°C. The appearing colonies were counted the next day.

### Fluorescence imaging

*B. anthracis* BYF10124 was injected into silkworm hemolymph. The silkworms were kept at 27°C. After 3h and 6h post-infection, hemolymph from the infected silkworm was collected, placed on a glass slide, and covered by a coverslip. Fluorescence images of the samples were collected using an inverted Zeiss LSM 780 confocal microscope equipped with an EM-CDD camera (Zeiss Research Microscopy Solutions, Germany) under a 40 x objective lens. To determine the effect of the antibiotic, 1 mg/kg ampicillin was injected into the infected larvae, and hemolymph was observed under the microscope.

### Assessment of virulence in silkworms

Virulence of the bacterial strains in silkworms was tested by injecting the exponentially growing *B. anthracis* 34F2 and mutants with disruption in virulence genes (~5 x 10^2^ CFU) into the hemolymph of each larva. For survival, larvae were observed at different time intervals post-infection. To determine microbial burden, hemolymph of the infected larva was collected at 3h and 6h post-infection, diluted in saline, and colony-forming units were determined.

## Results

### Silkworms are killed by *Bacillus anthracis* infection

To establish a silkworm infection model of *B. anthracis*, we injected silkworm larvae with different cell numbers of *B. anthracis* Sterne strain 34F2 into the hemolymph and checked the survival. We found that *B. anthracis* killed the silkworms in a dosedependent manner (**Fig. 1a**). We determined the lethal dose that killed 50% of the worms (LD_50_) 16h post-infection to be 8.3 x 10^2^ colony forming units (CFU) per larva (**Fig. 1b**). At 19h post-infection, when all the silkworms infected with 8.1 x 10^2^ CFU of *B. anthracis* died (**Fig. 1c**), saline-injected silkworms were still surviving (**Fig. 1d**), indicating that the fatality is brought about by *B. anthracis* infection. The death of larvae was accompanied by a change of skin color to black due to melanization.

**Figure 1:**
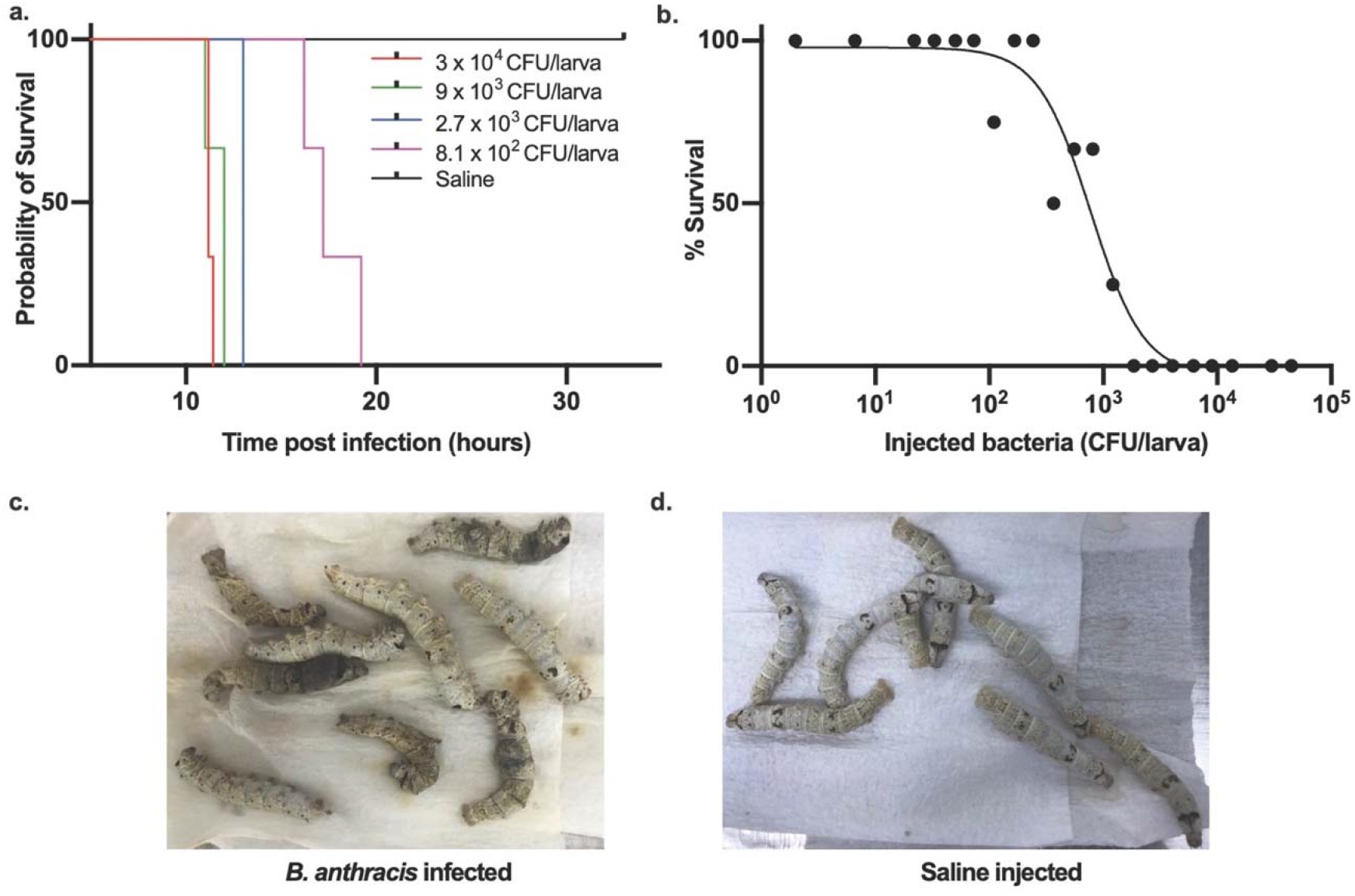
*Bacillus anthracis* kills silkworms. **a.** Dose-dependent killing of silkworms by *B. anthracis*. Data is representative of three independent experiments. **b**. Survival of silkworms 16h post-infection. Data is presented as a combined result of three independent experiments. LD_50_ is calculated by logistic regression analysis using the logit link function. **c.** Dead silkworms infected with *B. anthracis* (8.1 x 10^2^ CFU/larva) 19h post-infection. The dead silkworms turn black due to melanization. **d.** Alive silkworms injected with saline 19h post-injection.

To further confirm that the observed killing of silkworms was due to *B. anthracis* infection, we heat-killed the bacteria by autoclaving and injected into the silkworms. We found that injection of heat-killed bacteria equivalent to 1.5 x 10^6^ CFU did not kill the worms while live 2.6 x 10^3^ CFU killed the worms within 16h post-infection (**Fig. 2a**). We further constructed fluorescence protein AmCyan1-expressing *B. anthracis*, whose silkworm killing ability was similar to that of the wild-type (**Fig. 2b**). We confirmed the florescence-protein expression by observing the cells under a fluorescence microscope (**Fig. 2c**). We then infected the silkworm with the bacteria, recovered hemolymph 3h and 6h post-infection, and confirmed the fluorescence expression of bacteria in the hemolymph. While most of the bacteria were engulfed by hemocytes 3h post-infection (**Fig. 2d**), increased bacterial growth outside the hemocytes was observed 6h post-infection (**Fig. 2e**), indicating the progression of bacterial proliferation within the host and establishment of infection by the bacteria.

**Figure 2:**
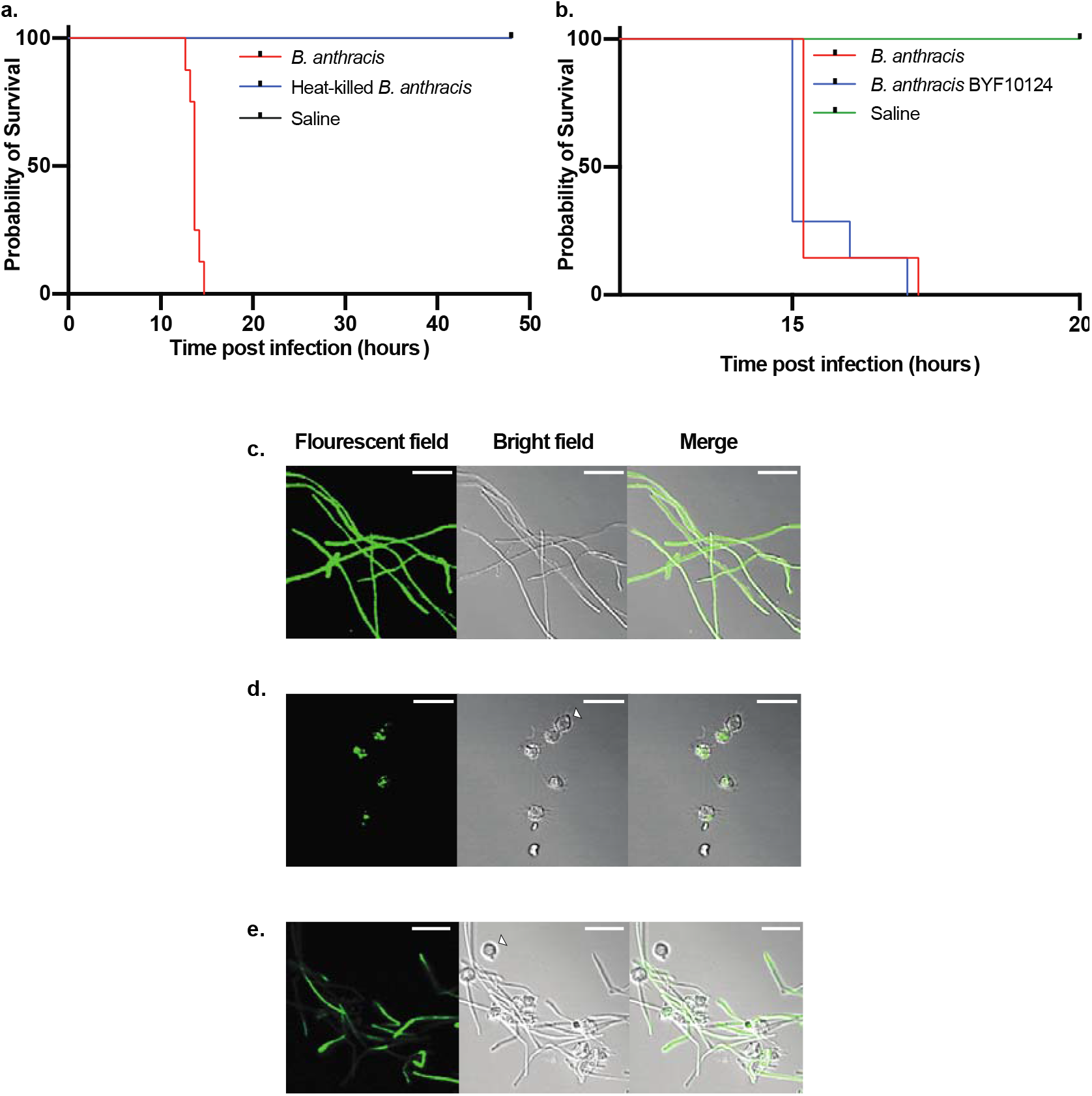
Live bacteria is required for silkworm killing. **a**. Survival of silkworms after injection of heat-killed *B. anthracis*. Live *B. anthracis* (2.6 x 10^3^ CFU/larva) and heat-killed *B. anthracis* (1.5 x 10^6^ CFU/larva) were injected to silkworms (n=8), and survival was observed. Representative data of two independent experiments are shown. **b.** Survival of silkworms after infection with wild-type and BYF10124. Wild-type *B. anthracis* (5 x 10^2^ CFU/larva) and BYF10124 (8.5 x 10^2^ CFU/larva) were injected to silkworms (n=7), and survival was observed. Representative data of two independent experiments are shown **c.** *In vitro* grown BYF10124 under microscope. **d, e.** BYF10124 in silkworm hemolymph 3h (**d**) and 6h (**e**) post-infection under microscope. White arrows indicate hemocytes of silkworm hemolymph. Scale bars, 20 μm.

### Infection is cured by antibiotics treatment

After confirming that *B. anthracis* establishes infection in silkworm and kills them, we evaluated the therapeutic effectiveness of clinically used antibiotics against *B. anthracis* infection. At first, we determined the *in vitro* antimicrobial susceptibility of *B. anthracis* toward a range of antibiotics. Consistent with reported studies^42–44^, we found that it was susceptible to most of the antibiotics tested and resistant to bacitracin and fosfomycin (**Table 2**).

**Table 2:**
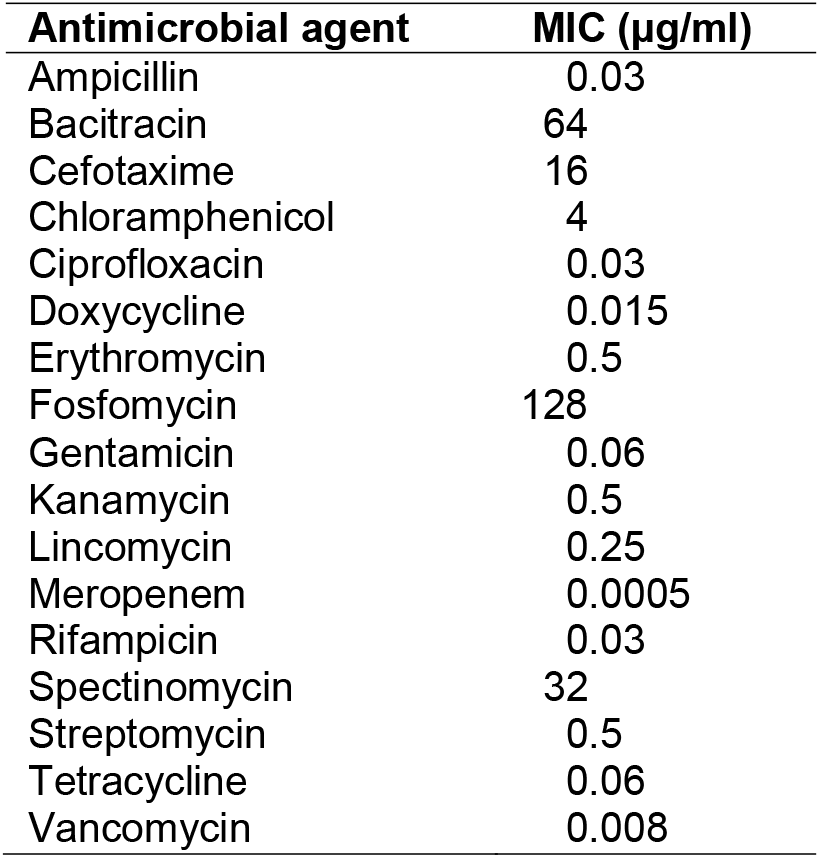
Antimicrobial susceptibility of *Bacillus anthracis*. MIC: minimum inhibitory concentration. Representative data of three independent experiments are shown.

Next, we selected two antibiotics that are commonly used for the treatment of anthrax, doxycycline and ampicillin, and injected them into the hemolymph of silkworms infected with *B. anthracis*. We found that both antibiotics cured silkworms and prevented their death (**Fig. 3a**). We further determined the dose-response of doxycycline and ampicillin and calculated the effective doses that cure 50% of the worms (ED_50_) 16h post-infection to be 0.05 mg/kg and 0.02 mg/kg, respectively (**Fig. 3b, c**). We, next, determined the bacterial burden in the silkworm hemolymph at different intervals of time post-infection and found that the number of viable cells decreased with time in the antibiotic-treated groups (**Fig. 3d**). In addition, we checked the fluorescence expression after administering ampicillin to the BYF10124 infected larvae. We found that after 3h post-infection, most of the hemocytes were colonized with bacteria (**Fig. 4a**). While upon ampicillin treatment, only a few hemocytes were colonized with bacteria, and the overall presence of bacteria was decreased (**Fig. 4b**). At 6h post-infection, bacteria started proliferating outside the hemocytes (**Fig. 4c**), while the ampicillin treated group had fewer bacteria engulfed in the hemocytes with no bacterial growth outside the hemocytes (**Fig. 4d).**

**Figure 3:**
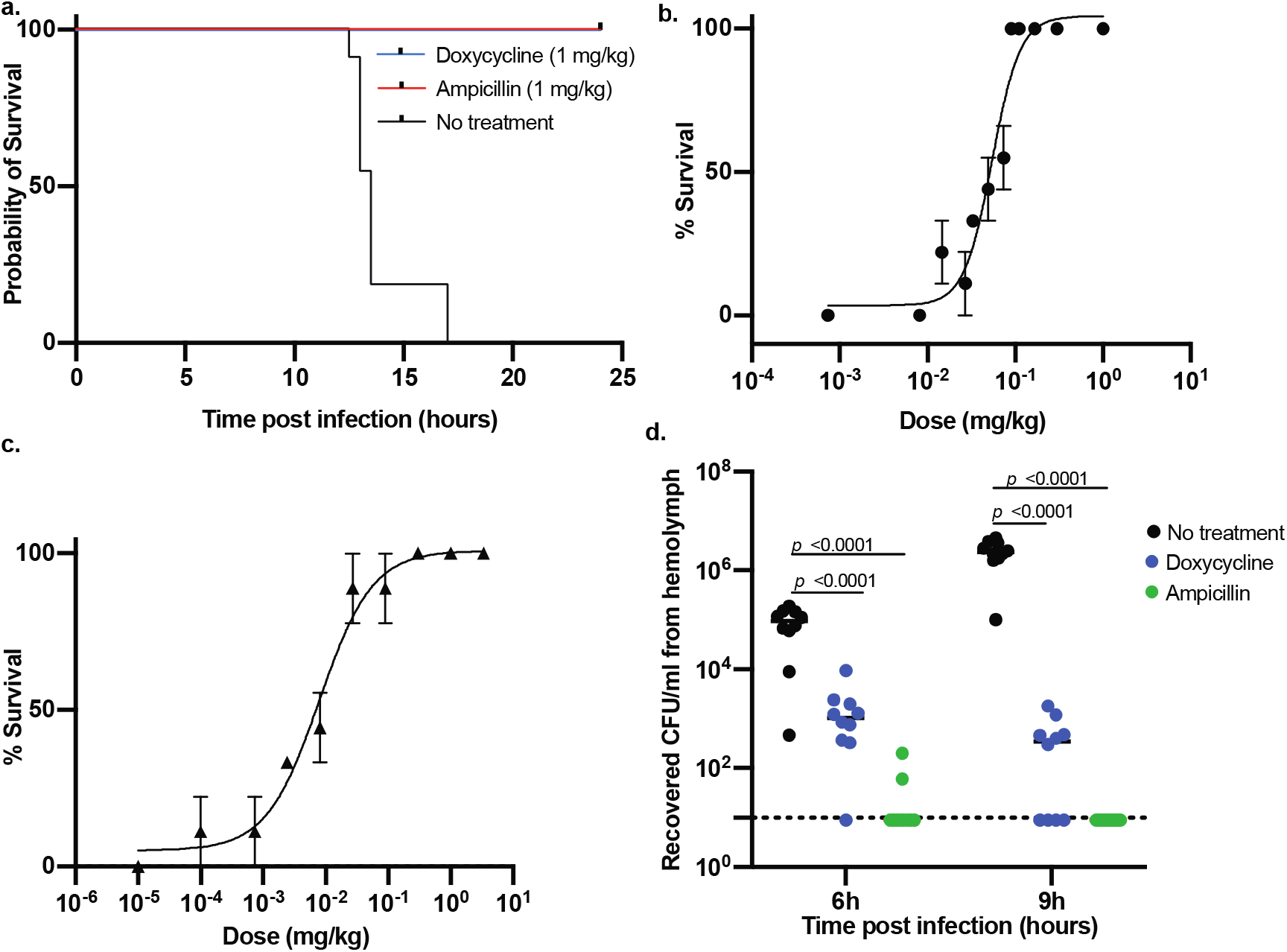
Treatment of *B. anthracis* infection by antibiotics. **a.** Survival of silkworms (n=10) with and without antibiotics treatment. Representative data from three independent experiments are shown. **b, c.** Survival of silkworms treated with various concentrations of doxycycline (**b**) or ampicillin (**c**) 16h postinfection. Data are mean ± SEM of three independent experiments. ED_50_ value was calculated by logistic regression analysis using the logit link function. **d.** Bacterial burden after 6h and 9h post-infection with and without antibiotics treatment (1 mg/kg). Statistical analysis was performed by one-way analysis of variance (ANOVA) with Dunnett’s multiple comparison test compared with the wild-type. Dotted line shows limit of detection.

**Figure 4:**
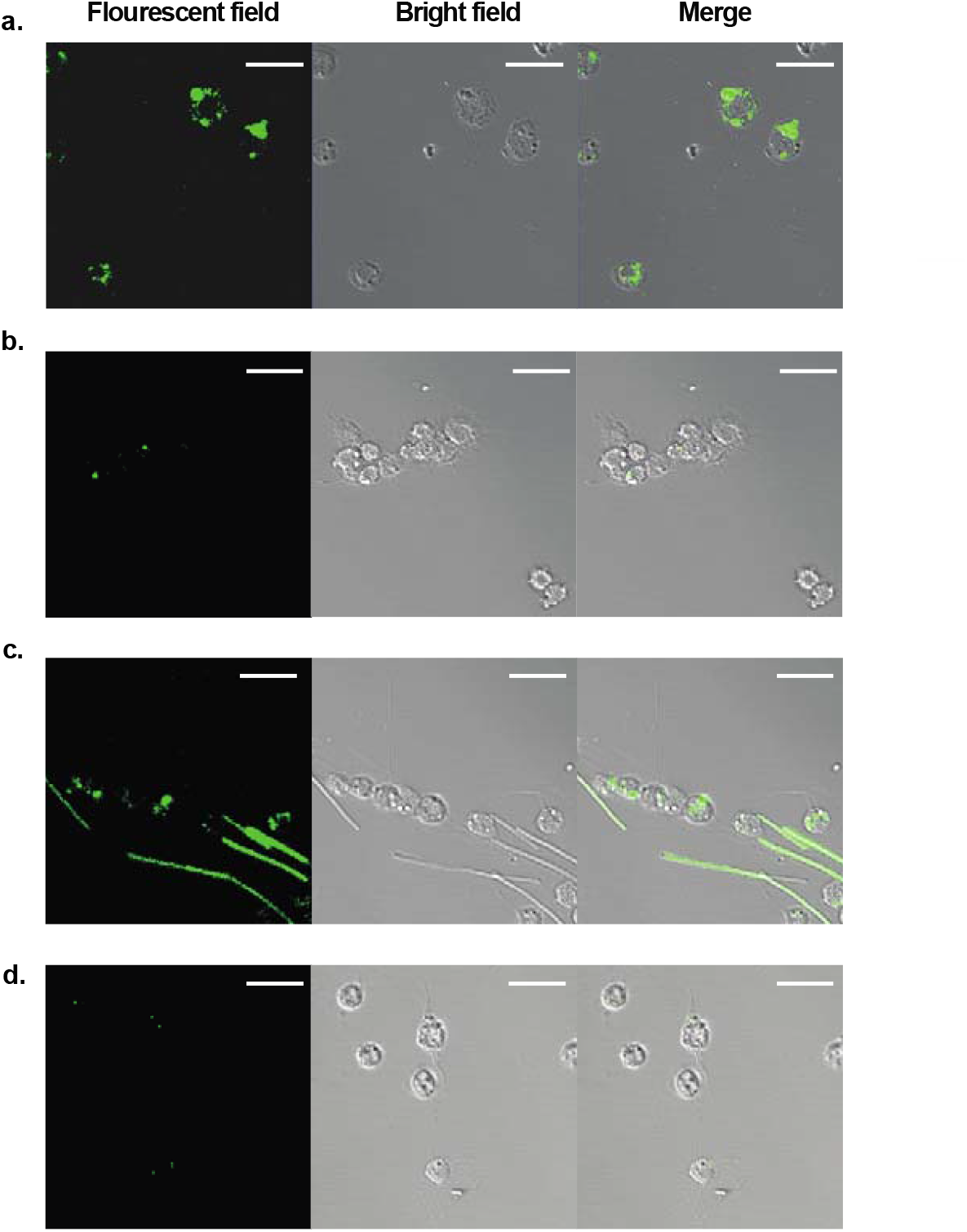
Clearance of *B. anthracis* from silkworm hemolymph upon antibiotic treatment. Time course of *B. anthracis* clearance by antibiotic treatment at 3 h (**a, b**) and 6 h (**c, d**) post-infection is shown. Silkworms were infected with BYF10124 and injected with either vehicle (**a, c**) or 1 mg/kg ampicillin (**b, d**). At specified time, hemolymph was obtained and visualized under the microscope. Scale bars, 20 μm.

### Silkworm as a model to assess virulence of *B. anthracis*

With the establishment of the silkworm infection model of *B. anthracis* as shown above, we next used silkworms to evaluate the virulence of *B. anthracis* mutants. The toxin-related genes are known to have roles in *B. anthracis* virulence^45,46^. To test whether these toxins also act on silkworms, we used mutants with disruptions in *pagA, lef*, and *atxA* ^41^. Located within a pathogenicity island on pXO1^11^, the *pagA, lef*, and *atxA* genes code for the protective antigen, the lethal factor, and a global virulence regulator AtxA, respectively. AtxA is reported to, directly and indirectly, regulate the transcription of several genes, including the *pagA* and *lef* genes^41,47^. We found that these mutants were less virulent in silkworms as they took a longer time to kill the larvae (**Fig. 5a**) and had attenuated colonizing ability (**Fig. 5b**). Taken together, it was evident that the disruption of virulence-related genes decreases the virulence of *B. anthracis* to the silkworm.

**Figure 5:**
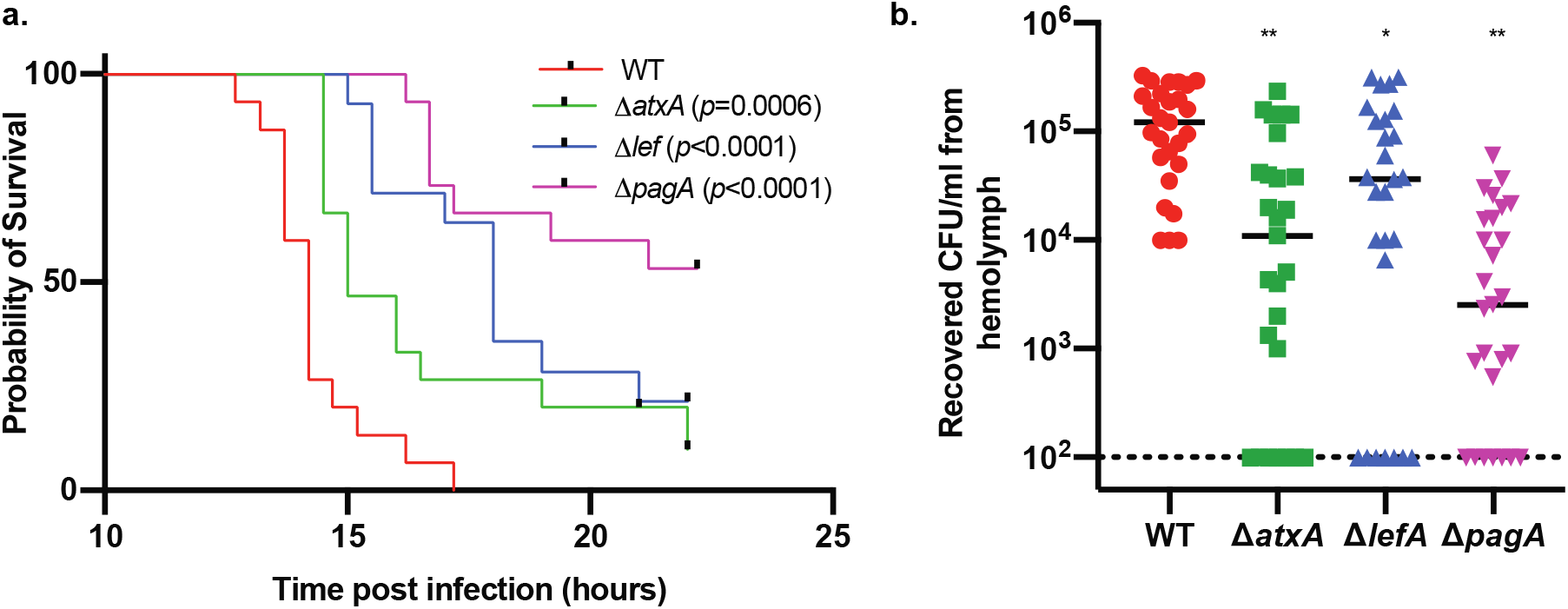
Assessment of virulence of *B. anthracis* mutants using silkworms. **a.** Survival of silkworms after infection with wild-type and virulence gene-disrupted mutants. Exponentially growing bacteria were injected into the hemolymph of silkworms and survival was observed. Data are shown as combined result from two independent experiments (n=15). Injected CFU/larva: WT (wild-type)= 5 x 10^2^ and 2.6 x 10^3^; Δ*atxA* = 4 x 10^2^ and 3.4 x 10^3^; Δl*ef*= 7.5 x 10^2^ and 2 x 10^3^; Δ*pagA* = 4.4 x 10^3^ and 1.0 x 10^4^. Statistical analysis was performed by Mantel-Cox log-rank test. **b**. Microbial burden of *B. anthracis* wild-type and mutants in silkworms 6h post-infection. Exponentially growing bacteria were injected into the hemolymph of silkworms, hemolymph was recovered 6h post-infection and CFU was determined. Data are shown as combined result from two independent experiments (n=27). Injected CFU/larva: WT (wild-type)= 2 x 10^2^ and 3 x 10^2^; Δ*atxA* = 1.8 x 10^2^ and 4.4 x 10^2^; Δ*lef*= 2.2 x 10^2^ and 2.4 x 10^2^; Δ*pagA* = 1.1 x 10^2^ and 2.8 x 10^2^. Statistical analysis was performed by one-way analysis of variance (ANOVA) with Dunnett’s multiple comparison test compared with the WT (* p < 0.05, ** p <0.0001). Dotted line represents limit of detection.

## Discussion

In this study, we established a silkworm model of *B. anthracis* infection and evaluated the therapeutic effects of clinically used antibiotics in silkworms infected with *B. anthracis*. Moreover, we generated a *B. anthracis* strain expressing AmCyan1, which was useful in evaluating the proliferation of bacteria inside the host over time. While silkworm infection models of human pathogens have been reported^30,32,48–50^, this is the first report of the silkworm infection model of *B. anthracis*. We found that *B. anthracis* Sterne killed silkworms and the LD_50_ was 8.3 x 10^2^ CFU, which was comparable with those in mice models where LD_50_ ranged from 1.6 x 10^2^ – 1.1 x 10^3^ CFU^51,52^. Live bacteria were required for silkworm killing, which was evident from the fact that heat-killed bacteria (10^6^ CFU) could not kill the larvae. *B. anthracis* established infection within silkworms as their proliferation was increased over time observed both from the increased CFU in the hemolymph of larvae and the increased proliferation of fluorescent bacteria under the microscope harvested at various intervals post-infection.

When we administered clinically used antibiotics to the *B. anthracis* infected silkworms in this study, we observed therapeutic activities of the antibiotics as they prolonged the survival of infected silkworms. Recovered CFU of bacteria from the treated silkworm hemolymph showed a faster clearance of bacteria in the ampicillin-treated group than in the doxycycline-treated group. As ampicillin is a bactericidal antibiotic, bacteria are killed in addition to clearance from the host immunity, which may have led to faster overall clearance; whereas, being a bacteriostatic antibiotic, doxycycline inhibited the bacteria growth, and overall clearance may have depended upon the host immunity taking a longer time. We further demonstrated, using fluorescent protein-expressing *B. anthracis*, that antibiotic treatment reduces bacterial burden in the hemolymph. The therapeutic effects of known antibiotics in the silkworm infection model imply that the therapeutic effectiveness of unknown compounds can be evaluated using this system, selecting for compounds with therapeutic activity and appropriate pharmacokinetics at an early stage of screening^35,36,38^. An additional advantage of using silkworms is that a small quantity of compounds would be enough to evaluate therapeutic effectiveness.

The innate immunity is partly conserved among silkworms and mammals, and several signaling cascades such as the mitogen-activated protein kinase (MAPK) pathways are activated in silkworms by bacterial components resulting in antimicrobial peptides production^53^. Thus, silkworms could differentiate the virulence of the mutant deficient in the lethal factor of *B. anthracis*, which acts via the MAPK pathway^54^. However, silkworms do not have acquired immunity, and therefore, they can be used to determine virulence factors that trigger the innate immunity and not the acquired immunity. Nonetheless, innate immunity is the first line of defense in all organisms^55^, and evaluation of virulence factors of pathogenic microorganisms have been performed using silkworm model^28,30,56–58^. The finding of this study showing attenuated virulence of strains of *B. anthracis* with disruption in known virulence genes suggested that silkworms can be used to evaluate the roles of unknown genes in the virulence of *B. anthracis*.

Since Sterne strain lacks pXO2 and is less virulent to higher animals^40^, silkworm model of *B. anthracis* Sterne infection will have an additional advantage for the identification of virulence factors encoded by genes either in the chromosome or pXO1 that might be masked in a highly virulent strain containing both pXO1 and pXO2. The identified virulence factors can then be utilized as potential vaccine targets in Sterne strain such as *htrA, aro, nos, sod*, and *clpX*^14,59–62^. As the use of Sterne strain in humans is associated with safety concerns, the identification of other virulence factors may contribute to the development of a safe human vaccine strain.

## Acknowledgments

We thank Dr. Takeshi Saito (International Institute for Zoonosis Control, Hokkaido University, Japan) for help with fluorescent microscopy and Tomoko Shimizu (International Institute for Zoonosis Control, Hokkaido University, Japan) for technical assistance.

## Funding

This work was supported by JSPS KAKENHI Grant Number JP19K16653 and JP21K15430 to AP, JP18K14672 to YF, and JP18K19436 to HH and AMED Grant number JP21wm0125008 (the Japan Program for Infectious Diseases Research and Infrastructure) to AP, YF, and HH.

## Declaration of interest statement

The authors declare no conflict of interest.

## Notes

### Competing Interest Statement

The authors have declared no competing interest.

